# Transposable elements impact the population divergence of rice blast fungus *Magnaporthe oryzae*

**DOI:** 10.1101/2023.05.12.540556

**Authors:** Lianyu Lin, Ting Sun, Jiayuan Guo, Lili Lin, Meilian Chen, Zhe Wang, Jiandong Bao, Justice Norvienyeku, Dongmei Zhang, Yijuan Han, Guodong Lu, Christopher Rensing, Huakun Zheng, Zhenhui Zhong, Zonghua Wang

## Abstract

Dynamic transposition of transposable elements (TEs) in fungal pathogens have significant impact on genome stability, gene expression, and virulence to the host. In *Magnaporthe oryzae*, genome plasticity resulting from TE insertion is a major driving force leading to the rapid evolution and diversification of this fungus. Despite their importance in *M. oryzae* population evolution and divergence, our understanding of TEs in this context remains limited. Here we conducted a genome-wide analysis of TE transposition dynamics in the 11 most abundant TE families in *M. oryzae* populations. Our results show that these TEs have specifically expanded in recently isolated *M. oryzae* rice populations, with the presence/absence polymorphism of TE insertions highly concordant with population divergence on Geng/*Japonica* and Xian/*Indica* rice cultivars. Notably, the genes targeted by clade-specific TEs showed clade-specific expression patterns and are involved in the pathogenic process, suggesting a transcriptional regulation of TEs on targeted genes. Our study provides a comprehensive analysis of TEs in *M. oryzae* populations and demonstrates a crucial role of recent TE bursts in adaptive evolution and diversification of the *M. oryzae* rice-infecting lineage.

**IMPORTANCE:** *M. oryzae* is the causal agent of the destructive blast disease, which caused massive loss of yield annually worldwide. The fungus diverged into distinct clades during adaptation toward two rice subspecies, Xian/indica and Geng/japonica. Although the role of TEs in the adaptive evolution was well established, mechanisms underlying how TEs promote the population divergence of *M. oryzae* remains largely unknown. In this study, we reported that TEs shape the population divergence of *M. oryzae* by differentially regulating gene expression between Xian/*Indica*-infecting and Geng/*Japonica*-infecting populations. Our results revealed a TE insertion mediated gene expression adaption that led to the divergence of *M. oryzae* population infecting different rice subspecies.

## INTRODUCTION

Rice blast disease, caused by the ascomycete filamentous fungus *Magnaporthe oryzae* (syn: *Pyricularia oryzae*), poses a significant threat to rice production worldwide, resulting in annual yield losses of 10-30% (1, 2). The deployment of resistant rice varieties is the most cost-effective and environmentally friendly strategy for controlling rice blast disease. However, the effectiveness of such resistance can rapidly be diminished due to rapid mutation accumulation in avirulence genes (3, 4). Therefore, it is crucial to uncover the mechanisms by which *M. oryzae* rapidly evolves and evades the rice immune system.

Based on the two-speed genome model, filamentous fungi genomes had been shown to display a bipartite architecture comprising of a gene-rich compartment, which evolves slowly containing core genes encoding essential functions and metabolisms, and a repeat-rich compartment, which evolves rapidly containing important virulence effectors involved in pathogenicity (5–10). To avoid recognition by the plant immune system, some pathogen effectors such as avirulence genes were shown to be influenced or silenced by transposon elements (9, 11). Transposable elements (TEs) make up over 10% of the *M. oryzae* genome (12, 13). Recent evidence has revealed that TEs can be inserted in or around important effectors and alter the virulence spectrum of *M. oryzae*. For example, the insertion of POT3 into the promoter region of AVR-Pita in *M. oryzae* led to the acquisition of virulence towards the resistant rice cultivar Yashiro-mochi (14). TEs are also able to affect gene expression networks, and TE-dependent transcriptional regulation of some essential effectors can facilitate the pathogen’s transition in its life cycle. For instance, in *Phytophthora sojae*, the avirulence genes *PsAvr1a*, *1b*, and *3a/5* were found to be transcriptionally inactive due to TE insertions in their promoter or 3’ UTR regions (15).

These studies have demonstrated the crucial role of TE-mediated genomic variations in pathogen adaptation. However, previous investigations have mainly focused on the impact of TEs on specific effectors or secreted proteins (13, 16–20). To gain a better understanding on the functions of TEs in the complex *M. oryzae*-rice pathosystem, we conducted a comprehensive analysis of TE insertion polymorphisms in 275 *M. oryzae* isolates (176 rice isolates and 99 non-rice isolates; **Table S1**), and systematically investigated the roles of TEs in the regulation of gene expression and population divergence in the *M. oryzae* rice population.

## RESULTS

### Recent large-scale TE bursts in the *M. oryzae* genome

To assess the activity of transposable elements (TEs) on the M. oryzae genome, we estimated the insertion time of long terminal repeat (LTR)-retrotransposons (LTR-RTs) by measuring the genetic distance between their 5’ and 3’ LTRs, which were identical at the time of TE insertion and gradually accumulated mutations over time. Using a de novo method that is based on the structure of LTR-RTs, we identified 1,129 intact LTR-RTs in seven near chromosomal-level *M. oryzae* rice isolates, including 70-15, Guy11, FJ81278, FJ98099, FJ72ZC7-77, AV1-1-1, and Sar-2-20-1 (12, 13, 21). Surprisingly, 91.7% (1,036/1,129) of the LTR-RTs showed extremely low levels of divergence between their LTR pairs (over 99% identity), and 69.4% (784/1,129) of them possessed identical LTR pairs, indicating that these LTR-RTs were recently inserted and could be still active in the *M. oryzae* genomes.

Then, we analyzed 11 TE families that are most abundant in the genomes of *M. oryzae* rice isolates, including six LTR-RTs (RETRO5, RETRO6, RETRO7, Maggy, MGLR-3, and Pyret), two non-LTR retrotransposons (MGL and Mg-SINE), and three DNA transposons (POT2, POT3, and Occan) (**Fig S1**) (12, 22–25). To assess the activity of these TE families, we calculated the Kimura 2-Parameter genetic distance (k-value) to measure the divergence between TE sequences and their associated consensus sequences (26). Low k-values indicate that the TE fragments were generated through recent insertion events, while high k-values indicate that the TE fragments are divergent copies generated through ancient transposition events (27). Our analysis revealed that all 11 TE families, especially sequences of MGL, Mg-SINE, Maggy, and a subset of POT2, POT3, and Occan, exhibit very low k-values, and more than half of all TE contents consisted of newly emerged TEs (k-values less than 5) (**Fig. 1a and b**), indicating a recent, large-scale burst of TEs in the genome of *M. oryzae* rice isolates. Notably, we observed two or more k-value peaks in POT2, Mg-SINE, and Pyret, indicating that these TEs families have undergone multiple rounds of amplification.

**FIG 1.**
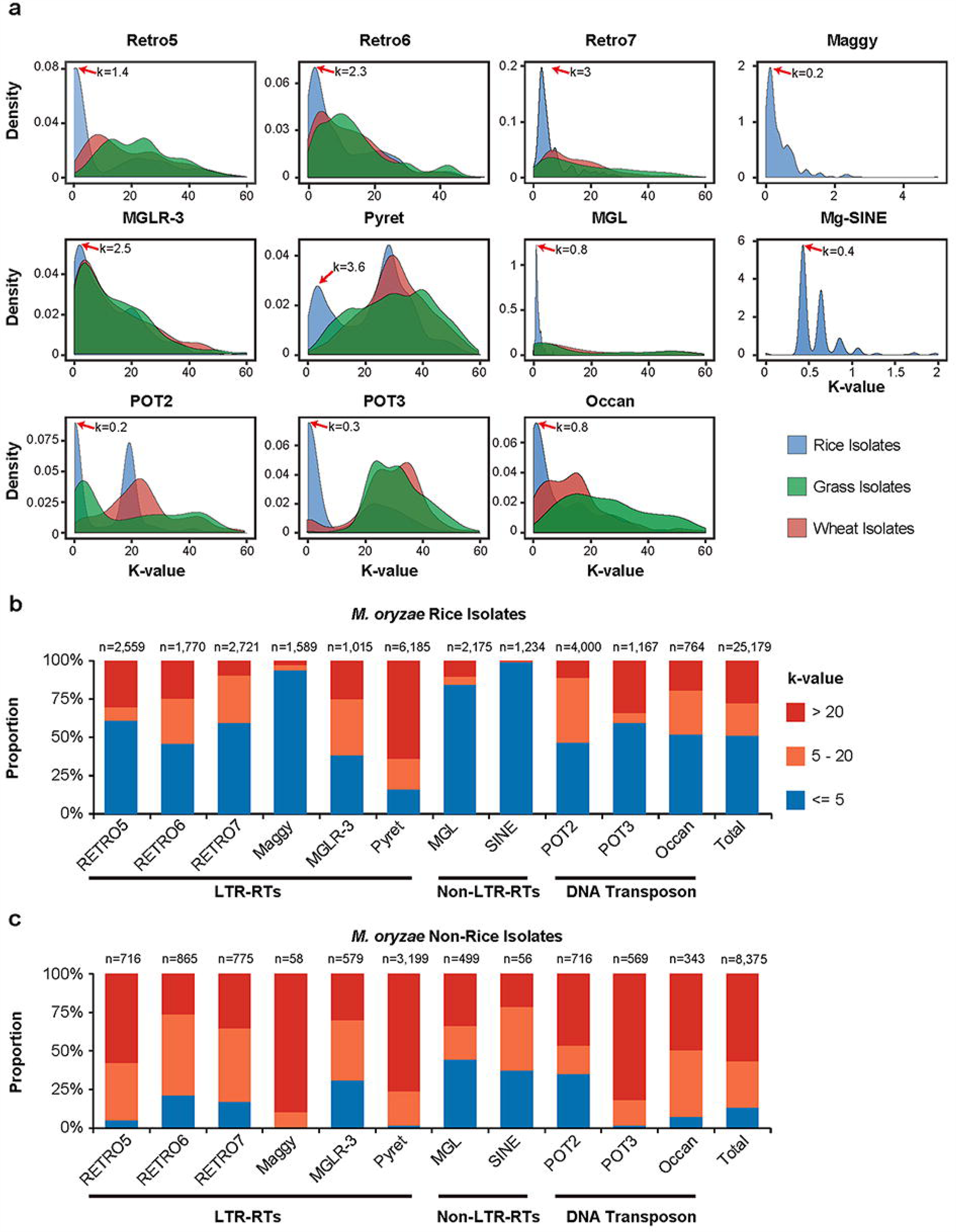
The distribution of 11 TE families in the *M. oryzae* population. **a)** The Kimura 2-Parameter genetic distance of 11 TE families in three different *Magnaporthe* species were shown in density plot. **b, c**) The k-value proportions of 11 TE families in *M. oryzae* rice isolates (**b**) and non-rice isolates (**c**). The numbers of TE fragment were marked at the top of bars.

Furthermore, we investigated these TE families in 275 genomes of *M. oryzae* isolates (**Table S1**) (28, 29). We found that TEs only accounted for less than five percent of the genomes of non-rice isolates, which is dramatically lower than that found in *M. oryzae* rice isolates (**Table S1 and S2**). Interestingly, the k-values of TEs were much larger and the proportion of newly emerged TEs was also much lower in *M. oryzae* non-rice isolates, indicating that the TEs were generated by more ancient insertion events and were inactive in *M. oryzae* non-rice isolates (**Fig. 1b and c**). Notably, the *M. oryzae* non-rice isolates contained only a few copies of fragmented Maggy and Mg-SINE, which were very abundant and possessed very low k-values in *M. oryzae* rice isolates, suggesting that the two TEs specifically amplified in *M. oryzae* rice isolates. In summary, our analysis demonstrated that the TEs were recently and specifically expanded in the genome of *M. oryzae* rice isolates and maintained high activity.

### Whole genome landscape of TE dynamics in *M. oryzae* population

To examine the dynamics of transposable elements (TEs) in the *M. oryzae* rice population, we conducted a genome-wide analysis of TE insertion sites in 90 rice isolates that had previously been published (30), with two *M. oryzae Setaria viridis* isolates as an outgroup (31). Using paired-end read mapping to the reference genome method, we identified a total of 11,163 TE insertion sites, with an average of 1,312 sites per isolate. To verify these insertion sites, 17 insertion sites randomly selected from Guy11 or FJ81278 isolates were proved to be presented as predicted through PCR-based genotyping or PacBio (**Table S3,S4**). The number (1,312 versus 739) and location of TE insertions differed dramatically between the rice and *S. viridis* isolates (**Fig. S2**), reflecting the evolutionary divergence of these subspecies and their corresponding TEs, which is consistent with the finding that these two subspecies diverged ~10,000 years ago (31). More than half (6,040/11,163) of the TE insertion sites were singletons specific to individual isolates (**Fig. 2**), indicating frequent TE transposition events. Notably, the number of POT2 insertion sites was substantially higher than those of other TE families, suggesting a higher activity and variability of POT2 in the *M. oryzae* rice population.

**FIG 2.**
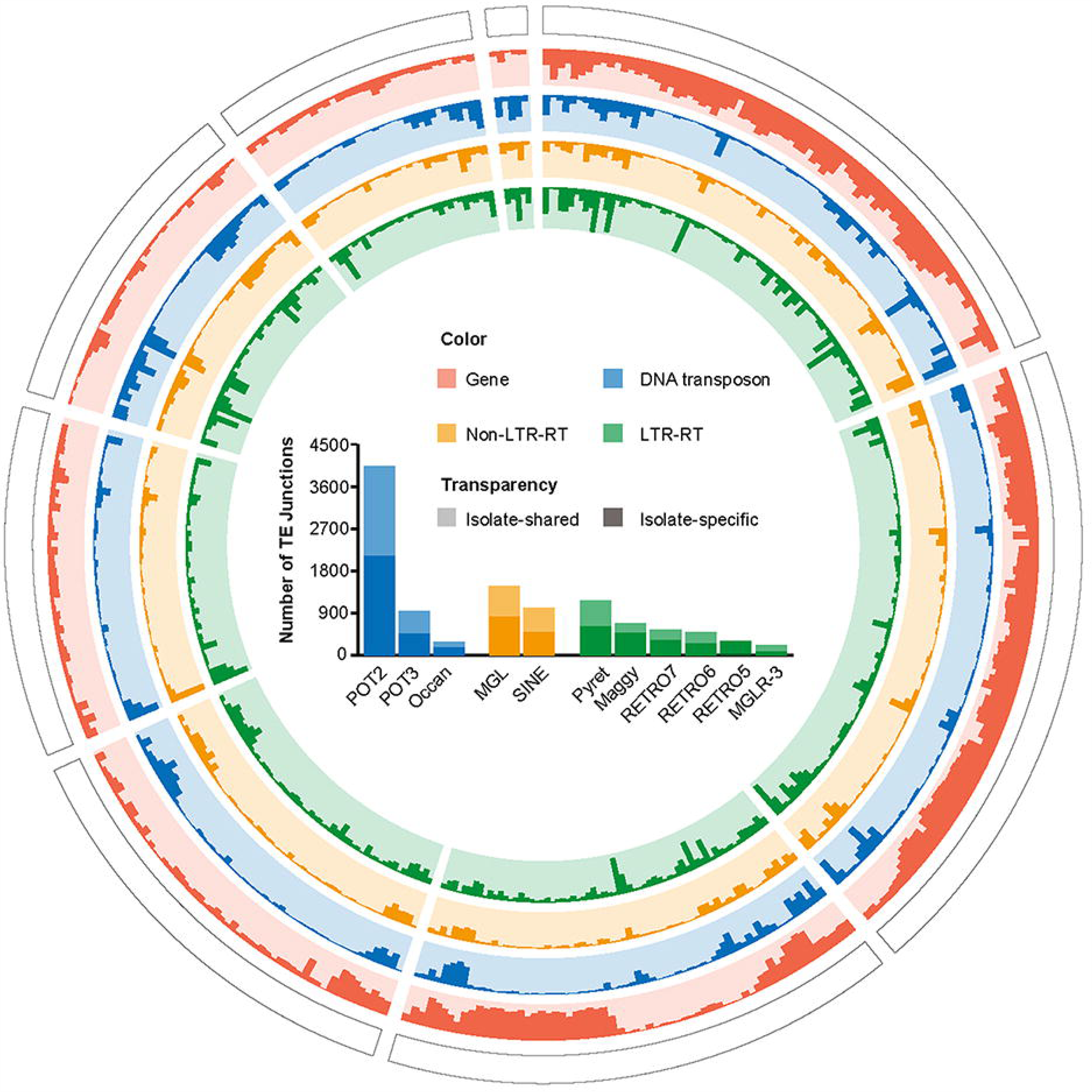
The number of TE junctions in the *M. oryzae* rice population. The circos diagram (from outer to inner) displays chromosomes, gene distribution, distribution of DNA transposon, Non-LTR-RT and LTR-RT junctions. The histogram shows the insertion site numbers of different TE families. The non-transparent and transparent colors represent the numbers of isolate-specific and isolate-shared TE junctions, respectively

Furthermore, we conducted a comprehensive analysis of the genomic distribution of TE insertion sites in the *M. oryzae* rice population. Of the 11,163 TE insertion sites identified, 77% (8,582/11,163) was found to be located within 1 kb of the flanking regions of genes or intragenic regions, and over 40% of the genes (5,259/12,991) were embedded by these TE insertions. Our enrichment analysis showed that the distribution of the 11 TE families is non-random in the *M. oryzae* rice isolates, and each family displayed a distinct preference for specific genomic regions. For instance, Maggy, MGLR-3, RETRO5, RETRO7, Pyret, POT3, and Occan were predominantly distributed in intergenic regions, while POT2 displayed a preference for the gene flanking regions. Additionally, SINE, MGL, and RETRO6 were found to preferentially target intragenic regions (adjusted p-value < 0.01, **Table S5**). These findings provide valuable insights into the mechanisms underlying TE insertions and their potential impact on gene regulation in *M. oryzae*.

### Higher frequency of POT2 and POT3 insertions in promoter of secreted proteins

We observed that genes encoding secreted proteins were more closely associated with TE junctions (**Fig. 3a**), and the proportion of genes encoding secreted proteins with TE insertion within 1-kb flanking regions was significantly (p=7.6e-4) higher than in those of non-secreted proteins (**Table 1**). Moreover, enrichment analysis for TEs associated with genes encoding secreted proteins showed that POT2 and POT3 were overrepresented in promoters of genes encoding secreted protein (adjusted p-value < 0.01, **Table S6**), implying that genes encoding secreted proteins are more prone to disruption by POT2 and POT3.

**FIG 3.**
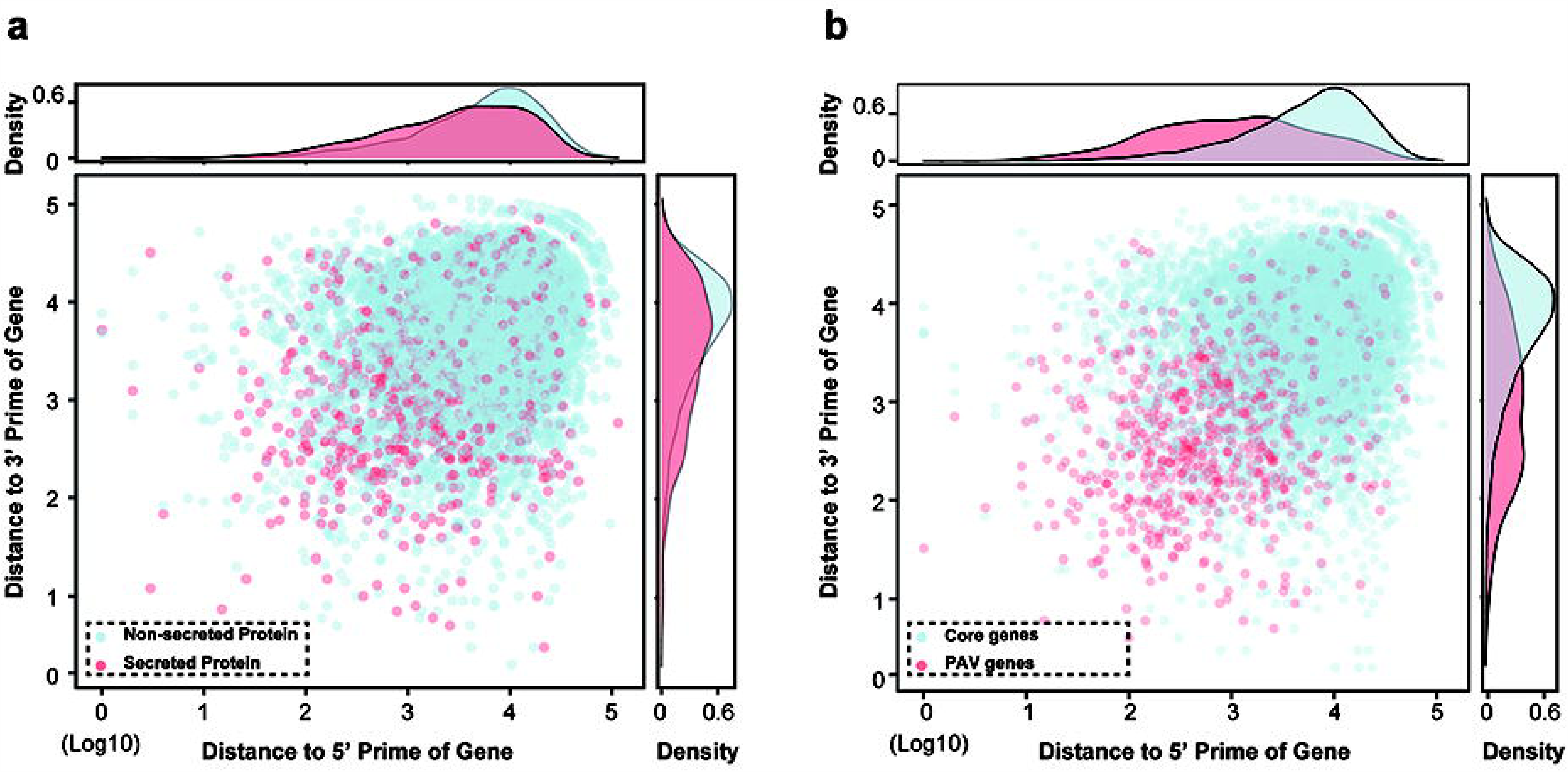
The TEs were in proximity to secreted proteins and PAV genes. The distance of TEs to the 5’ and 3’ ends of secreted proteins and non-secreted proteins (**a**) or core genes and PAV genes (**b**).

Previous studies have shown that genes with presence/absence variation (PAV) tended to be located near transposable elements (TEs) in fungal pathogen genomes, while core genes were located further away from the TE-rich compartments (32–34). In this study, we compared the genomic distribution of core genes and PAV genes and found that PAV genes tended to be located closer to TE insertion sites than core genes (**Fig. 3b**). Our results were consistent with previous findings and suggested that PAV genes may be more susceptible to TE-mediated disruption, potentially contributing to their faster evolution in the context of host-pathogen interactions. Together, these findings suggested that TEs play a role in the evolution of pathogen effectors and contribute to the dynamic nature of host-pathogen interactions.

### Association of TEs with *M. oryzae* rice population divergence

Considering that the *M. oryzae* rice population diverged within only one thousand years (30) and that the large-scale TE burst happened recently, we thus raised the question of whether the population divergence of *M. oryzae* is associated with recent TE amplification events. To characterize correlations between the 90 isolates, we estimated the distance for each of two isolates by calculating the identity of the TE insertion sites. The pairwise TE insertion identities varied from 17.6% between YN126441 and FJ12JN-084-3 to 87.2% between TW-PT3-1 and TW-PT6-1, with an average of only 38.7%, thereby strongly implicating the high variability of TE junctions between the different isolates. However, when we grouped these isolates based on the pairwise TE insertion identities, we discovered two distinct clusters (**Fig. 4a**) matching the Clade2 and Clade3 isolates that we had previously defined based on genome-wide SNPs (30). We noticed that the remaining isolates out of the two clusters were also able to match the Clade1 isolates even though they showed a relatively low pairwise TE insertion identity, which can be attributed to an earlier time of divergence of Clade1 isolates from the *M. oryzae* rice population when compared to the other two clade isolates. Furthermore, we found that the pairwise TE insertion identity between intra-clade isolates was higher than that between inter-clade isolates (**Table S7**). Surprisingly, the hierarchical tree constructed using TE insertion sites showed a high degree of similarity to the phylogenetic tree constructed based on whole-genome SNPs (**Fig. 4b**) (30), indicating that the recent TE amplification event has been a major force driving population divergence of *M. oryzae*. Consistently, principal component analysis of these TE insertion sites displayed a similar pattern (**Fig. 4c**). Considering that the TEs were largely amplified recently in the *M. oryzae* genome (**Fig. 1**), we presumed that the recent high activity of TEs has been one of the major forces driving population divergence of *M. oryzae*.

**FIG 4.**
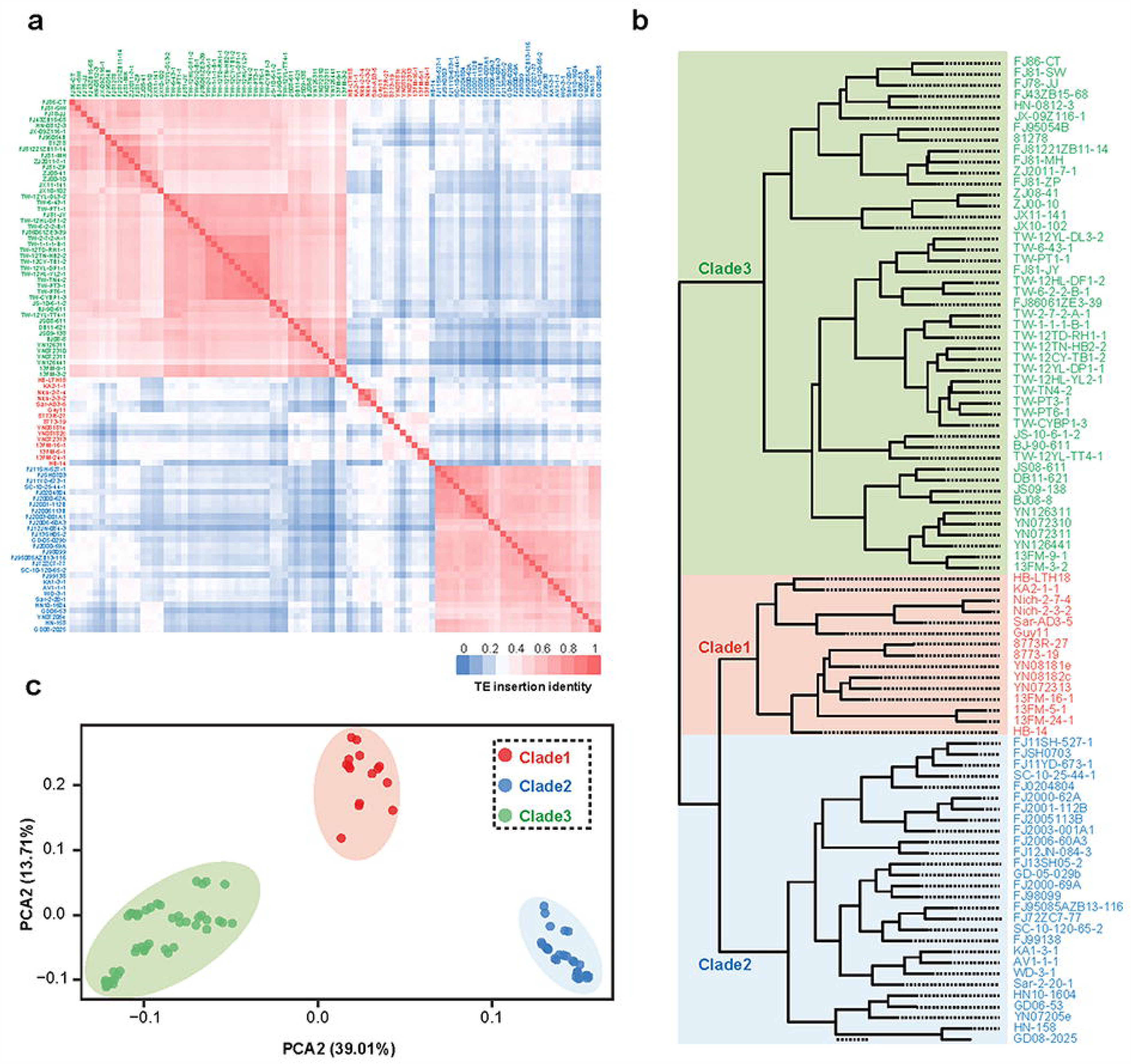
TE insertion was associated with the divergence of *M. oryzae* rice population. **a)** The heatmap showing the identity of TE insertion site between each two isolates. **b)** The hierarchical tree built by using the TE insertion sites of *M. oryzae* population. Three distinct clades were marked in red, blue and green colors respectively. **c)** Principal component analysis using the TE insertion sites.

### POT2 and Mg-SINE are critical for the divergence of Clade2 and Clade3 isolates

TEs are able to exert either beneficial or deleterious effects on host genomes, and the retention or elimination of TEs is largely determined by their impact on the host. Positive selection has been shown to drive the frequency of a TE locus to increase or decrease dramatically during a population bottleneck or in response to a new environment (35). We hypothesized that a portion of clade-specific TE insertion sites had been fixed in the intra-clade isolates, contributing to the adaptive evolution of the clade isolates. To identify the clade-specific TE insertion sites, we empirically screened out those that were present in at least 80% of the intra-clade isolates and absent in more than 80% of the other two clade isolates. A total of 11 Clade1-specific, 212 Clade2-specific, and 168 Clade3-specific TE insertion sites were identified, with the number of Clade1-specific TE insertion sites being too small for subsequent analysis. Enrichment analysis revealed that POT2 and Mg-SINE TE families were overrepresented in both Clade2 and Clade3 datasets, indicating that the retention of insertion of these two TE families in subpopulations of *M. oryzae* rice isolates was tightly associated with the divergence of the rice-infection population (**Fig. 5a and b**).

**FIG 5.**
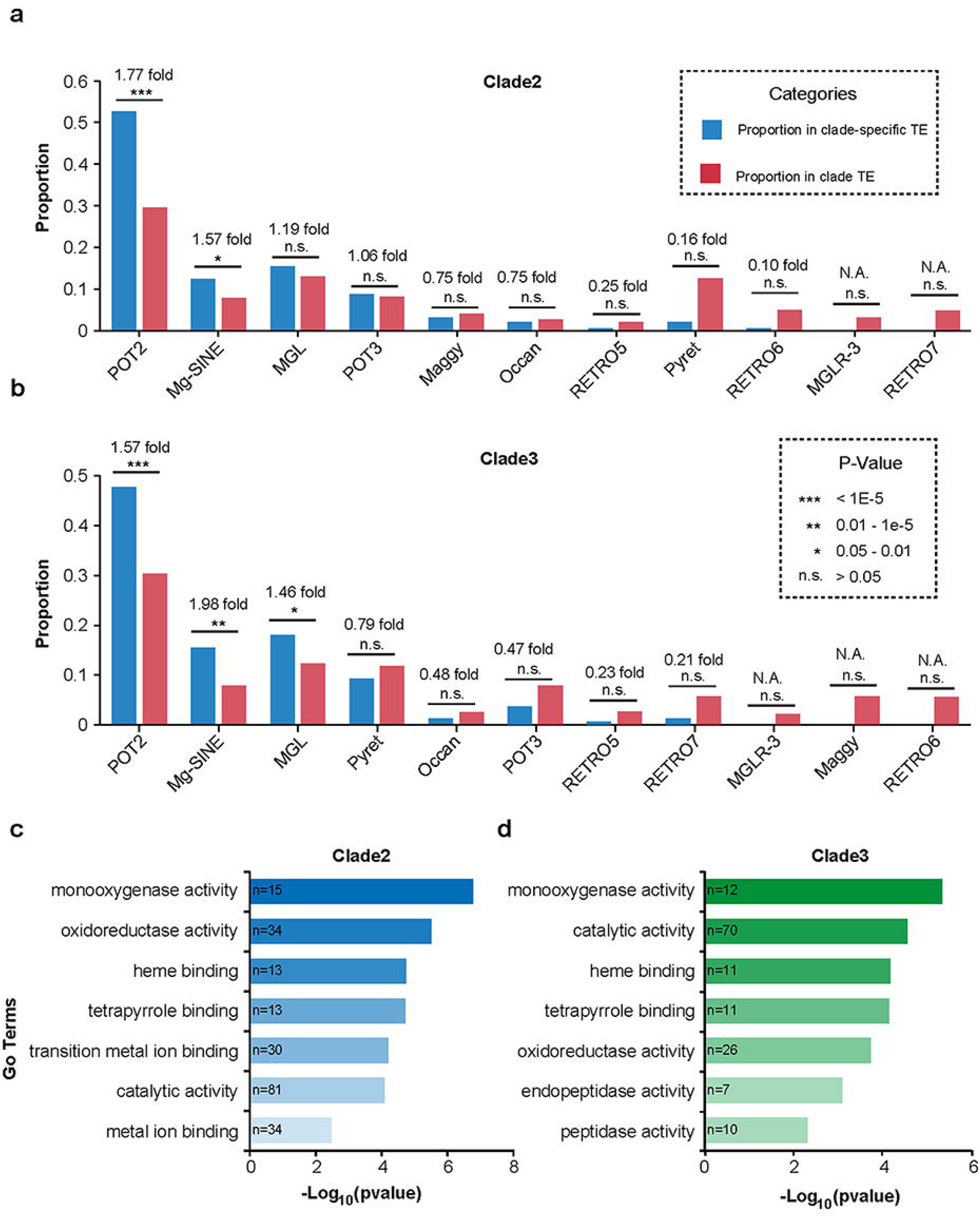
Characteristics of clade-specific TEs in the *M. oryzae* rice population. **a, b)** Enrichment analysis of 11 TE families in Clade2- (**a**) and Clade3-specific (**b**) insertion sites. Fisher’s exact test was used for significance test. **c**) GO enrichment analysis for the clade2- (**c**) and clade3-specific (**d**) TE-associated genes.

We investigated the influence of clade-specific TE junctions on genes, which, for this purpose, we referred to as <u>c</u>lade-<u>s</u>pecific <u>T</u>E-associated genes (*CSTs*). We identified a total of 238 Clade2-specific and 173 Clade3-specific TE-associated genes. Interestingly, less than 10% of these genes overlapped, suggesting that the Clade2- and Clade3-specific TEs have distinct targeted genes (**Fig. S2**). Gene ontology (GO) enrichment analysis revealed that the Clade2- and Clade3-specific TE-associated genes were enriched under similar GO terms (adjusted p-value < 0.01, **Fig. 5c, d**). Of note, the top enriched term in both datasets was ‘GO:0004497, monooxygenase activity,’ which is correlated to cytochrome P450s on the fungal genome. We further validated the GO enrichment results by performing enrichment analysis for these genes using the Pfam database (**Table S8**). Several previous studies have shown that fungal P450s possess detoxifying functions towards compounds produced by host plants during pathogen infection, thereby enhancing the fitness of the pathogenic fungus to specific host genotypes (31, 36–40). Therefore, we suggest that TE-induced variations in different members of P450s partially contribute to the differential pathogenicity of the two clade isolates.

### Genes associated with TE are significantly lower expressed

Previous studies have demonstrated that transposable elements (TEs) were able to affect gene expression by inserting into gene promoters or intragenic regions (41–43). Therefore, we investigated whether TE insertion polymorphisms could shape the gene expression networks in *M. oryzae* rice populations. To this end, we selected 16 isolates and extracted total RNA for sequencing (**Table S9**). We identified 2,282 genes targeted by 4,236 TE insertion sites that were polymorphic between the 16 isolates. We then grouped the genes based on whether they contained TE insertion sites (TE-present or TE-absent) and compared the expression levels between these two groups. We observed that the TE-present gene group displayed significantly (p=2.17e-26) lower expression levels than the TE-absent gene group (**Fig. 6a**), suggesting that TEs have negative effects on the expression of their target genes. We have identified 131 genes that exhibited clade-specific expression patterns. Furthermore, based on the expression levels of these genes, we were able to divide 16 isolates into three distinct clusters using principal component analysis. These findings suggest that transposable element-mediated gene regulation had a profound impact on population divergence (**Fig. 6b and c**). Interestingly, we found that only genes with insertion polymorphisms of POT2 and Mg-SINE displayed transcriptional suppression, while other TE families displayed little impact on the expression of their target genes. Our results suggest that the insertion polymorphisms of POT2 and Mg-SINE were able to shape the gene expression network of *M. oryzae* by inducing transcriptional suppression.

**FIG 6.**
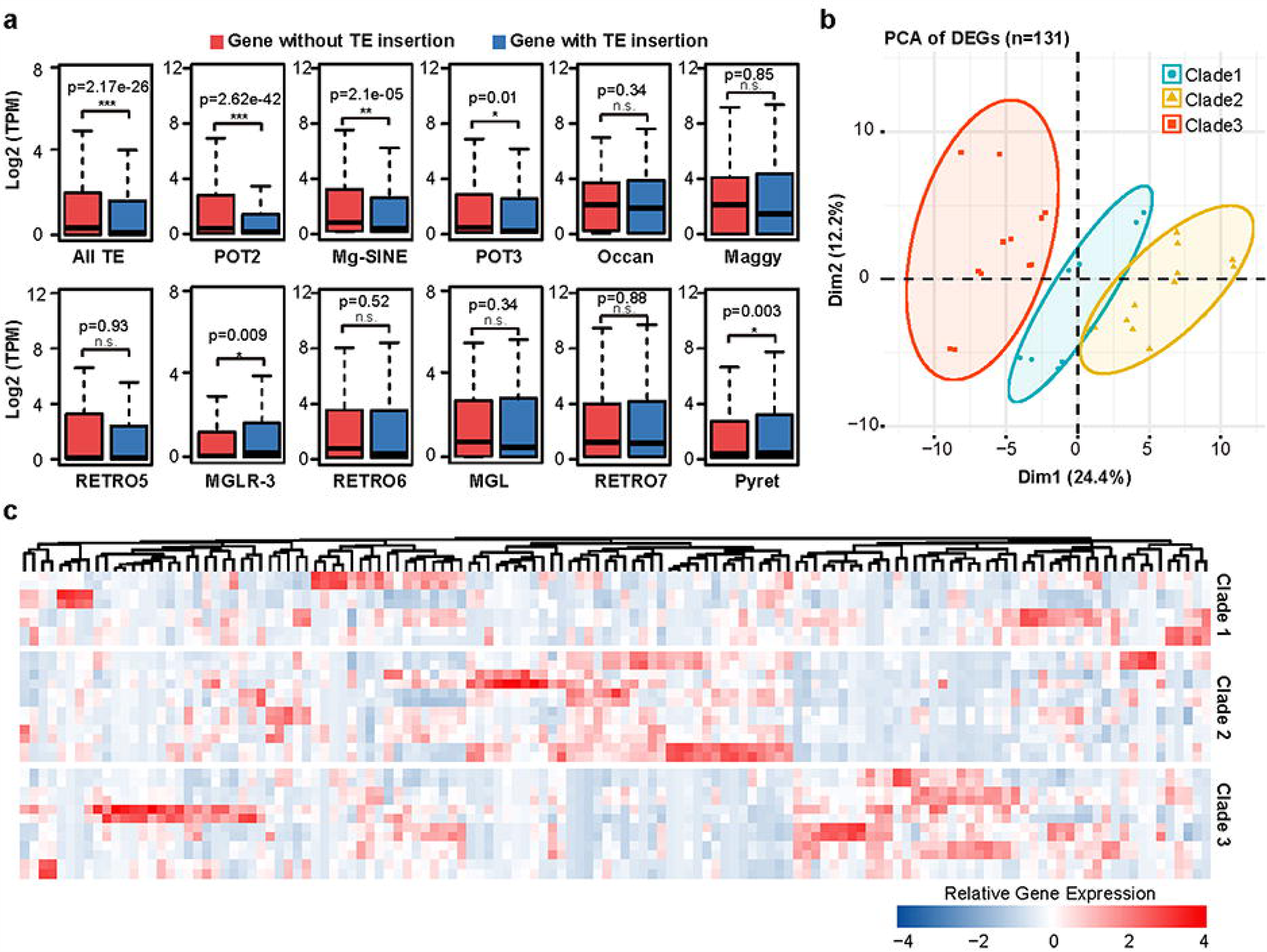
The impact of clade-specific TE insertion on gene expression. **a)** Comparison of gene expressions between the isolates present or absent with TE insertion. **b)** PCA clustering of 16 isolates based on expression level of 131 clade-specific DEGs. **c)** Heatmap showing expression level of 131 clade-specific DEGs in 16 isolates.

### *CST6* and *CST10* were required for the pathogenicity of *M. oryzae*

To further investigate the impact of clade-specific TE in the virulence divergence within *M. oryzae* rice population, we selected 15 of the clade-specific TE-associated genes (CSTs) for functional analyses (**Fig. 7a and Table S10**). We defined CSTs as genes that have TE insertion exclusively present in one clade and absent in another clade. Among them, CST1-7 are Clade1-specific, and are only transcriptionally active in Clade 2 isolates, while CST8-15 are Clade2-specific, and are only transcriptionally active in Clade 1 isolates. We amplified CST1-7 from a Clade 2 isolate, 95085, and ectopically overexpressed them in a Clade 1 isolate FJ81278. Conversely, CST8-15 were amplified from a Clade 1 isolate, FJ81278, and ectopically overexpressed in a Clade 2 isolate 95085 (CST8-11) or transiently expressed in tobacco leaves (CST12-15). The functional analyses revealed a role of CST6 and CST10 in the pathogenicity of *M. oryzae* by leaf punch inoculation assays on the two Japonica cultivars (NPB and TP309) and the two Indica cultivar (CO39 and MH63) (**Fig. 7b and c**). While the CST6-OE strain produced smaller lesions and reduced fungal biomass compared with the wild type, suggesting a potential role as a negative regulator of virulence (**Fig. 7b and c**). The CST10-OE strain was more aggressive and produced larger lesions and increased fungal biomass compared with the wild type, suggesting its function as a positive regulator of virulence (**Fig. 7d and e**). These results suggest that CST can profoundly impact virulence in *M. oryzae*, influencing its interaction with different rice species.

**FIG 7.**
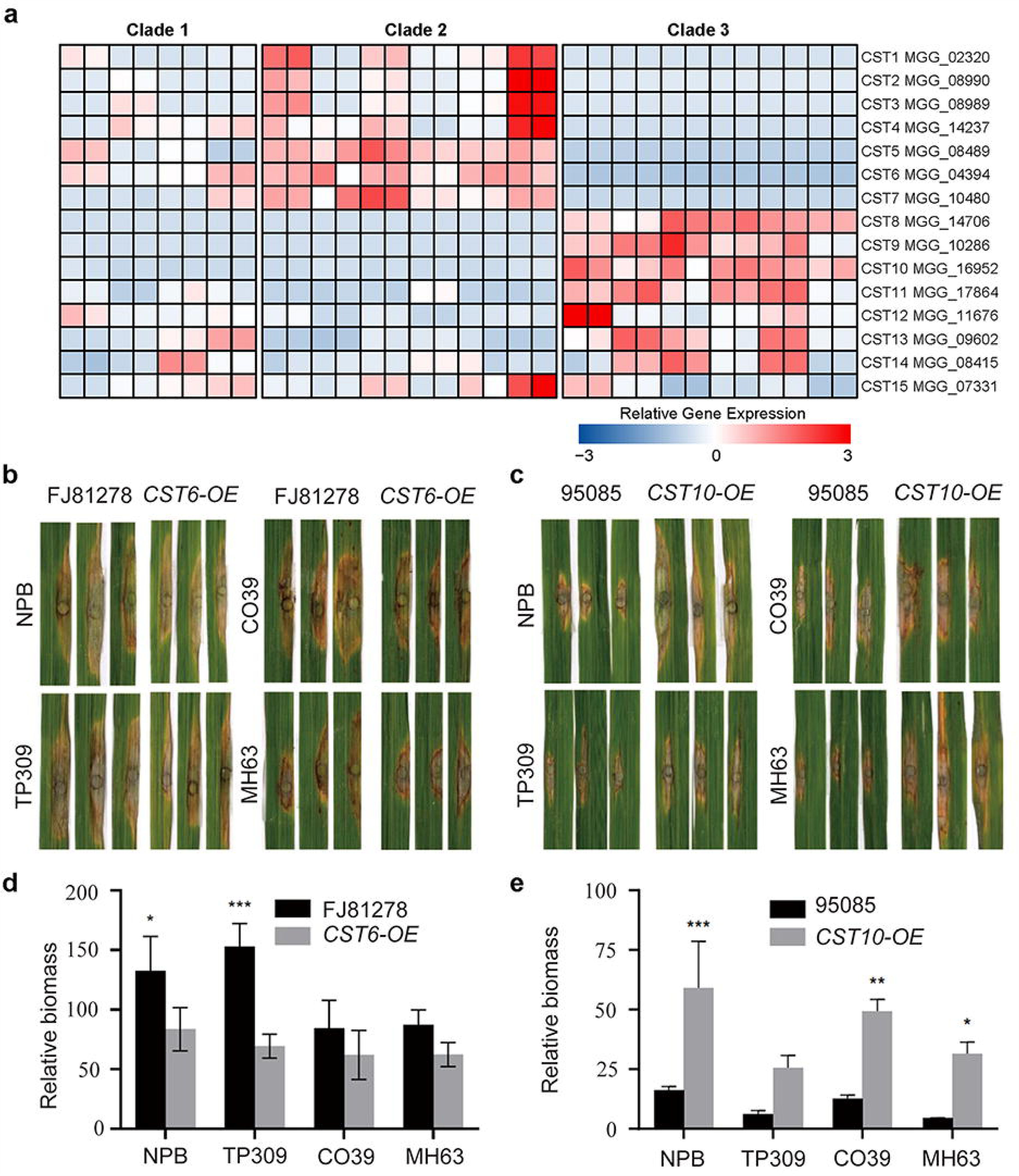
CST6 and CST10 were required for the pathogenicity of *M. oryzae* rice isolates. **a)** Heatmap showing expression level of 15 clade-specific TE-associated genes (CSTs) in 16 isolates. **b)** Leaf punch inoculation assays were conducted to assess the impact of CST6 overexpression on two Japonica cultivars (NPB and TP309) and the two Indica cultivar (CO39 and MH63). The Clade3 isolate (FJ81278), lacking CST6 expression, was used as the wild-type control. **c)** Leaf punch inoculation assays were conducted to assess the impact of CST10 overexpression on two Japonica cultivars (NPB and TP309) and the two Indica cultivar (CO39 and MH63). The Clade3 isolate (95085), lacking CST10 expression, was used as the wild-type control. **d)** Relative fungal biomass on leaf punch inoculation assays of CST6 overexpression on two Japonica cultivars (NPB and TP309) and the two Indica cultivar (CO39 and MH63). **e)** Relative fungal biomass on leaf punch inoculation assays of CST10 overexpression on two Japonica cultivars (NPB and TP309) and the two Indica cultivar (CO39 and MH63).

## DISCUSSION

The high genomic plasticity and rapid evolution of the plant pathogen *M. oryzae* presents a severe challenge for rice blast disease control (44, 45). Previous studies have identified transposons as a major driving force for the adaptive evolution of fungal pathogen genomes (10). Insertion polymorphisms of transposable elements (TEs) have been shown to lead to genomic instability, increased chromosomal recombination, and accelerated gene evolution (17, 46). However, the precise role of TE dynamics during *M. oryzae* evolution has remained poorly understood. To address this, we conducted a population-scaled TE analysis of *M. oryzae* to investigate how TE insertion polymorphisms contributed in shaping population structure and divergence.

Previous study estimated that the *M. oryzae* rice population diverged 175 to 2,700 years ago (45). Here, we employed two methods to assess the activity of transposable elements (TEs) in the genome of *M. oryzae*. First, we estimated the insertion time of LTR-RTs and secondly, we calculated the Kimura 2-parameter genetic distance for the 11 most abundant TE families. Both analyses indicate that TEs have undergone recent amplification in the *M. oryzae* genome and have remained highly active. Previous studies have suggested that the insertion of TEs having caused gene presence/absence variations to be the main evolutionary mechanism driving the divergence of host-specific *M. oryzae* lineages (16). In this study, we investigated the abundance and activity of 11 TEs in pathogens from wheat and grass infecting lineages, and found that these 11 TEs have undergone specific expansion in *M. oryzae* rice isolates while remain inactive in non-rice isolates. Based on these results, we propose that the recent burst of TE activity could be the primary factor responsible for the divergence of *M. oryzae* from other host infecting lineages and rice subspecies.

To elucidate the role of TEs in genome evolution and divergence, we conducted a genome-wide survey of TE insertion sites in 90 *M. oryzae* rice isolates (30). We utilized PoPoolationTE2, which is a validated method for estimating TE insertion frequency in populations (47). The TE insertion junctions varied widely among the 90 isolates, indicating the highly dynamic and active feature of TEs in *M. oryzae* rice populations. Consistent with prior research, we found that the distribution of TE junctions across the genome was not evenly distributed, and different TE families or superfamilies displayed preferences for insertion into distinct genomic regions (7, 17, 19).

TEs have been shown to be a major contributor in causing genomic variations, and the evolution of plant pathogens (19, 48, 49). Previous studies have reported that in the major wheat pathogen *Zymoseptoria tritici*, recent TE bursts were associated with the proximity to genes (48). And TE insertion repressed the expression of *REP9-1* (49) or *Avr3D1* (50), and consequently resulted in the altered virulence in different isolates of this fungus. In the polyphagous fungal pathogen *Rhizoctonia solani*, TEs mediated the structural variations of regions encoding pathogenicity associated genes (51). Similarly, TE insertions in or around *Avr* genes in *M. oryzae* were able to lead to transcriptional silencing and loss of avirulence function (14, 52–56). Our analysis revealed that TE junctions are frequently observed in the flanking regions of genes encoding secreted proteins (SPs), and the proportion of genes encoding SPs with TE insertions is higher than that of non-SPs. This is consistent with previous findings that SPs are enriched in repeat-rich regions and are prone to rapid evolution (13, 57). We propose that the variation in SPs, induced by TE insertion polymorphisms, is able to facilitate adaptive evolution of *M. oryzae*. Presence/absence variations in genes resulting from TE insertions have been identified to be associated with the divergence of host-specific *M. oryzae* lineages (16). Consistent with this, we found that genes exhibiting high gain/loss polymorphisms in the 90 isolates were preferentially located near TE junctions, suggesting that TE-mediated presence/absence variation constitutes a significant mechanism underlying differentiation of host-specific or intra-species *M. oryzae* rice isolates.

Through the comparison of TE junctions in 90 *M. oryzae* rice isolates, we observed a clustering pattern that was similar to the three previously defined *M. oryzae* clades. This led us to investigate the potential role of TEs in *M. oryzae* rice population divergence. We constructed a hierarchical tree based on the TE junctions and found that it closely resembled the phylogenetic tree constructed using genome-wide SNPs. Principal component analysis of the TE junctions also yielded a similar clustering pattern. These findings suggested that the transposition of TEs was strongly associated with *M. oryzae* rice population divergence. Given that both the junction of the majority of TEs in the *M. oryzae* rice isolate genomes and the divergence of *M. oryzae* rice population were occurred recently, we hypothesize the recent burst of TEs to be a driving force contributing to the *M. oryzae* rice population divergence.

TE loci that undergo positive selection during evolution will be retained and will exhibit high frequencies in a population. Based on the hierarchical tree and PCA results, we postulate that the intra-clade isolates have a fixed set of TE insertion sites that are specific to each clade. These clade-specific TE insertion sites are then able to serve as molecular markers to distinguish isolates belonging to the three clades. Interestingly, we found that POT2 and Mg-SINE were enriched in clade-specific TE insertion sites, suggesting that these clade-specific insertions of POT2 and Mg-SINE were beneficial for the adaptive evolution of clade isolates. Furthermore, we noted that cytochrome P450 proteins were overrepresented in both Clade2- and Clade3-specific TE-targeted genes. Given the essential roles of P450s in detoxifying phytoalexins produced by host plants, we hypothesize that the differential pathogenicity of clade isolates may be partially due to the variations in P450s induced by clade-specific TE insertions.

TEs integrated within or flanking genes have been shown to induce gene silencing through the formation of heterochromatic regions (10, 43). To investigate the correlation between TE insertion and gene expression regulation in *M. oryzae*, we performed RNA sequencing on 16 isolates from the three clades and conducted functional studies on genes disrupted by clade-specific TEs. Our analysis revealed that genes containing a TE insertion displayed significantly lower expression compared to genes without TE insertion. Notably, only POT2 and Mg-SINE insertions led to substantial suppression of gene expression. The initial functional analysis suggests that these CSTs play a crucial role in the pathogenic process. Surprisingly, these CSTs exhibit dual functionality, acting as both negative and positive regulators of virulence, while the detailed mechanisms underlying these roles require further investigation. Therefore, we hypothesized that clade-specific insertions of POT2 and Mg-SINE contribute to the adaptive evolution of clade isolates by regulating expression of specific genes and affecting adaptive traits.

Through a comprehensive analysis of TEs in populations of the rice-infecting fungus *M. oryzae*, we have revealed the crucial roles played by recent TE insertional bursts in the adaptive evolution and diversification of this fungal lineage. We demonstrate that recent TE insertions have led to the emergence of genes with clade-specific expression patterns, contributing significantly to the divergence of *M. oryzae* rice population and their adaptation to different rice subspecies. These findings highlight the significance of TE-mediated genetic changes in the regulation of gene expression, which in turn contributes to clade divergence and allows *M. oryzae* to adapt to diverse environmental pressures, including those imposed by different rice cultivars. Our findings highlight the significance of TE-mediated genetic changes in the regulation of gene expression, which in turn contributes to clade divergence.

## MATERIALS AND METHODS

### RNA extraction, library generation, and sequencing

The fungal strains were cultured in liquid CM medium by incubation at 28°C under shaking at 110 rpm for three days. The mycelium was filtered, washed with double-deionized water, and dried before being ground in liquid nitrogen. Ground samples were transferred into DNase/RNase-free Eppendorf tubes, suspended in 1 mL Trizol, and vortexed vigorously. To eliminate proteins, 200 μL of chloroform was added to the mixture, which was then shaken for 15 s. After centrifugation at 12,000 rpm for 15 min at 4°C, 400 μL of the supernatant was collected and mixed with 400 μL of cold isopropanol. The mixture was kept at −20°C for at least two hours, then centrifuged at 12,000 rpm for 15 min at 4°C. The supernatant was discarded, and the precipitates were washed with 1 mL of 70% alcohol and centrifuged at 12,000 rpm for 5 min at 4°C. After air drying for 5 min at room temperature, the pellets were diluted with 54 μL of DNase/RNase-free deionized water and treated with 2 μL of DNase I at 37°C for 30 min. The mixture was brought up to 800 μL with RNase-free water, followed by the addition of an equal volume of chloroform. After gentle mixing, the mixture was centrifuged at 12,000 rpm for 15 min at 4°C. About 500 μL of the supernatant was collected and mixed with 500 μL of cold isopropanol. The mixture was then kept at −20°C for more than three hours, followed by centrifugation at 12,000 rpm for 15 min at 4°C. The precipitates were washed with 1 mL of 70% alcohol, air-dried for 5 min at room temperature, and eluted with DNase/RNase-free deionized water. The RNA samples were then stored at −80°C for RNA sequencing analysis.

### Estimation of TE activity

LTR-Finder (58) with modified parameters of ‘-D 15000 -d 1000 -L 7000 -l 100 -p 20 -C -M 0.8’, LTR-harvest (59) with modified parameters of ‘-similar 80 -vic 10 -seed 20 -seqids yes -minlenltr 100 -maxlenltr 7000 -mintsd 4 -maxtsd 6 -motif TGCA -motifmis 1’ and LTR-Retriever, and LTR-Retriever (60) with default parameter were used for *de novo* identification of full-length LTR-RTs and estimation of insertion time. Information on TE classification is based on previous research (12) and the conserved domains of TE consensus sequences were predicted by Conserved Domain Database (61). The Kimura 2-Parameter genetic distances (k-values) of TE fragments were calculated by RepeatMasker (62) with option ‘-a’.

### Prediction of secreted proteins and PAV genes

To identify putative secreted proteins, several criteria were employed, including the presence of a signal peptide cleavage site, the absence of a transmembrane domain, and a protein length of less than 400 amino acids. SignalP 4.0 and TMHMM 2.0 were utilized for signal peptide and transmembrane domain prediction, respectively (63, 64). The transcript sequences of the 70-15 reference genome were aligned to the previously published assemblies of 90 isolates using NCBI-blastn with default parameters. Genes that exhibited more than 90% similarity with no gaps longer than 50 bp when compared to the assemblies were marked as “present”. Genes that did not meet these criteria were marked as “absent”. PAV (presence-absence variation) genes were defined as those that were absent in more than five isolates, while core genes were defined as those present in all isolates.

### Identification of TE insertion sites in *M. oryzae* populations

*M. oryzae* 70-15 TE is annotated by RepeatMasker with 11 TEs as TE library. The pair-end reads were mapped to the *M. oryzae* 70-15 reference genome using BWA (65) with default parameters. The alignment files were exported to PoPoolation2 (66) with default parameter to identify TE insertion site for each isolate. The TE insertion sites with score less than 0.3 were filtered, and the insertion sites located within 50 bp were considered as one insertion event.

### Construction of TE hierarchical tree, principal component analysis

The insertion sites that are present in at least 5 isolates were used for constructing hierarchical tree and principal component analysis (PCA). The TE insertion sites in presence/absence variation format were exported to the R package ‘hclust’ (67) for construction of TE hierarchical tree. Vcftools (68) and Plink (69) were used for PCA.

### Functional enrichment analysis

GO annotation information of *M. oryzae* was obtained from the JGI database (70). Conserved domains of *M. oryzae* protein sequences were predicted using the Pfam database (71). Fisher right-tailed test was used for enrichment analysis and a cutoff p-value less than 0.05 was used to define significant enrichment.

### RNA-seq analysis

The clean reads were mapped to the 70-15 reference genome using hisat2 v2.2.1 (72) with default parameters. Stingtie v2.1.4 was used for expression quantification (73). Gene expressions were normalized with transcripts per million (TPM). The expression of each gene in the isolates with or without TE insertion were counted, averaged and compared. Two-tailed Wilcoxon paired test was used to estimate the significance of expression difference.

### Pathogenicity assay of the *CST*-overexpressing transformants

The coding sequence (CDS) of *CST6* and *CST10* were amplified from the Clade 2 isolate, 95085, and the Clade 1 isolate, FJ81278, respectively, and were inserted into the *pKNT-RP27* vector. The resulting constructs were then introduced into FJ81278 (*CST6*) or 95085 (*CST10*), respectively. The PEG-mediated protoplast transformation was performed as described previously (74). To determine the role of selected *CST* genes in the pathogenicity of *M. oryzae*, punch inoculation was performed as previously described (75). In brief, 10 μL spore solution (5×10^5^ spores/mL in sterilized water containing 0.02% Tween) was inoculated into wounded rice leaves. The inoculated rice plants were placed in a greenhouse. Disease symptoms were recorded 10 days post inoculation. DNA extracted from the diseased rice leaves was subjected to the quantitative fluorescence analyses. The primers used in this study were listed in Table S11.

## DATA AVAILABILITY

All high-throughput sequencing data generated in this study are accessible at NCBI’s Gene Expression Omnibus (GEO) via GEO Series accession number GSE205351 (https://www.ncbi.nlm.nih.gov/geo/query/acc.cgi?acc=GSE205351). The consensus TE sequnces of *M. oryzae* (https://github.com/S-t-ing/mBio-data-availablility/blob/ <u>main/Mo.TE_Consensus.fasta</u>), Genome-wide annotation of TEs in the genomes of 275 *Magnaporthe* isolates (https://github.com/S-t-ing/mBio-data-availablility/tree/ <u>main/gff</u>), and the TE insertion sites among the 90 *Magnaporthe* rice isolates (https://github.com/S-t-ing/mBio-data-availablility/blob/main/TE%20insertion%2090%20isolates.xlsx) were available online.

## ACKNOWLEDGMENTS

This work was financially supported by the National Natural Science Foundation of China (32272513 and 32172365) and Central Guidance on Local Science and Technology Development Fund of Fujian Province (2022L3088). Open Access funding provided by Fujian Agriculture and Forestry University.

---

**TABLE 1** The numbers of secreted protein and non-secreted protein that have TE insertion in 1-kb flanking regions or within gene.

## SUPPLEMENTAL MATERIAL

**FIG S1** Schematic of the 11 most abundant TE families on the genome of the *M. oryzae* rice isolate. The conserved domains in the relative location of the TE consensus were highlighted using different colors.

**FIG S2** The venn chart compares the TE insertion loci of 90 *M. oryzae* rice isolates and two *S. viridis* isolates.

**FIG S3** The venn chart displays the intersection of clade2-specific and clade3-specific TE-associated genes.

**TABLE S1** The proportions of 11 TE families on the genomes of 275 *M. oryzae* isolates.

**TABLE S2** The proportions of 11 TE families on the genomes of a *M. oryzae* wheat isolate and three grass isolates.

**TABLE S3** Validation of 17 TE insertion sites by PacBio.

**TABLE S4** Validation of 17 TE insertion sites by PCR amplification.

**TABLE S5** Insertion preference of various TEs in different genomic regions.

**TABLE S6** Enrichment analysis of TEs in the promoter of secreted proteins.

**TABLE S7** Identity of TE insertion sites between intra- or inter-clade isolates.

**TABLE S8** Enrichment analysis of the clade2- and clade3-specific TE-associated genes using conserved domains.

**TABLE S9** GEO information of 16 RNA seq samples.

**TABLE S10** Basic information about 15 clade-specific TE-associated genes (CSTs).

**TABLE S11** Primers used in this study.

## REFERENCES

1. Wilson RA, Talbot NJ. 2009. Under pressure: investigating the biology of plant infection by Magnaporthe oryzae. Nat Rev Microbiol 7:185–95.

2. Asibi AE, Chai Q, Coulter JA. 2019. Rice blast: A disease with implications for global food security. Agronomy 9:451.

3. Leach JE, Vera Cruz CM, Bai J, Leung H. 2001. Pathogen fitness penalty as a predictor of durability of disease resistance genes. Annu Rev Phytopathol 39:187–224.

4. Wang B-h, Ebbole DJ, Wang Z-h. 2017. The arms race between Magnaporthe oryzae and rice: Diversity and interaction of Avr and R genes. Journal of Integrative Agriculture 16:2746–2760.

5. Dong S, Raffaele S, Kamoun S. 2015. The two-speed genomes of filamentous pathogens: waltz with plants. Current opinion in genetics & development 35:57–65.

6. Chuong EB, Elde NC, Feschotte C. 2017. Regulatory activities of transposable elements: from conflicts to benefits. Nat Rev Genet 18:71–86.

7. Bourque G, Burns KH, Gehring M, Gorbunova V, Seluanov A, Hammell M, Imbeault M, Izsvák Z, Levin HL, Macfarlan TS. 2018. Ten things you should know about transposable elements. Genome biology 19:1–12.

8. Gout L, Kuhn ML, Vincenot L, Bernard-Samain S, Cattolico L, Barbetti M, Moreno-Rico O, Balesdent MH, Rouxel T. 2007. Genome structure impacts molecular evolution at the AvrLm1 avirulence locus of the plant pathogen Leptosphaeria maculans. Environ Microbiol 9:2978–92.

9. Raffaele S, Kamoun S. 2012. Genome evolution in filamentous plant pathogens: why bigger can be better. Nat Rev Microbiol 10:417–30.

10. Faino L, Seidl MF, Shi-Kunne X, Pauper M, van den Berg GC, Wittenberg AH, Thomma BP. 2016. Transposons passively and actively contribute to evolution of the two-speed genome of a fungal pathogen. Genome Res 26:1091–100.

11. Seidl MF, Thomma B. 2017. Transposable Elements Direct The Coevolution between Plants and Microbes. Trends Genet 33:842–851.

12. Dean RA, Talbot NJ, Ebbole DJ, Farman ML, Mitchell TK, Orbach MJ, Thon M, Kulkarni R, Xu JR, Pan H, Read ND, Lee YH, Carbone I, Brown D, Oh YY, Donofrio N, Jeong JS, Soanes DM, Djonovic S, Kolomiets E, Rehmeyer C, Li W, Harding M, Kim S, Lebrun MH, Bohnert H, Coughlan S, Butler J, Calvo S, Ma LJ, Nicol R, Purcell S, Nusbaum C, Galagan JE, Birren BW. 2005. The genome sequence of the rice blast fungus Magnaporthe grisea. Nature 434:980–6.

13. Bao J, Chen M, Zhong Z, Tang W, Lin L, Zhang X, Jiang H, Zhang D, Miao C, Tang H, Zhang J, Lu G, Ming R, Norvienyeku J, Wang B, Wang Z. 2017. PacBio Sequencing Reveals Transposable Elements as a Key Contributor to Genomic Plasticity and Virulence Variation in Magnaporthe oryzae. Mol Plant 10:1465–1468.

14. Kang S, Lebrun MH, Farrall L, Valent B. 2001. Gain of virulence caused by insertion of a Pot3 transposon in a Magnaporthe grisea avirulence gene. Mol Plant Microbe Interact 14:671–4.

15. Whisson S, Vetukuri R, Avrova A, Dixelius C. 2012. Can silencing of transposons contribute to variation in effector gene expression in Phytophthora infestans? Mobile genetic elements 2:110–114.

16. Yoshida K, Saunders DG, Mitsuoka C, Natsume S, Kosugi S, Saitoh H, Inoue Y, Chuma I, Tosa Y, Cano LM, Kamoun S, Terauchi R. 2016. Host specialization of the blast fungus Magnaporthe oryzae is associated with dynamic gain and loss of genes linked to transposable elements. BMC Genomics 17:370.

17. Thon MR, Pan H, Diener S, Papalas J, Taro A, Mitchell TK, Dean RA. 2006. The role of transposable element clusters in genome evolution and loss of synteny in the rice blast fungus Magnaporthe oryzae. Genome Biol 7:R16.

18. Castanera R, Lopez-Varas L, Borgognone A, LaButti K, Lapidus A, Schmutz J, Grimwood J, Perez G, Pisabarro AG, Grigoriev IV, Stajich JE, Ramirez L. 2016. Transposable Elements versus the Fungal Genome: Impact on Whole-Genome Architecture and Transcriptional Profiles. PLoS Genet 12:e1006108.

19. Muszewska A, Steczkiewicz K, Stepniewska-Dziubinska M, Ginalski K. 2019. Transposable elements contribute to fungal genes and impact fungal lifestyle. Sci Rep 9:4307.

20. Shirke MD, Mahesh HB, Gowda M. 2016. Genome-Wide Comparison of Magnaporthe Species Reveals a Host-Specific Pattern of Secretory Proteins and Transposable Elements. PLoS One 11:e0162458.

21. Zhong Z, Lin L, Zheng H, Bao J, Chen M, Zhang L, Tang W, Ebbole DJ, Wang Z. 2020. Emergence of a hybrid PKS-NRPS secondary metabolite cluster in a clonal population of the rice blast fungus Magnaporthe oryzae. Environ Microbiol doi:10.1111/1462-2920.14994.

22. Sanchez E, Asano K, Sone T. 2011. Characterization of Inago1 and Inago2 retrotransposons in Magnaporthe oryzae. Journal of general plant pathology 77:239–242.

23. Kachroo P, Leong SA, Chattoo BB. 1994. Pot2, an inverted repeat transposon from the rice blast fungus Magnaporthe grisea. Mol Gen Genet 245:339–48.

24. Farman ML, Tosa Y, Nitta N, Leong SA. 1996. MAGGY, a retrotransposon in the genome of the rice blast fungus Magnaporthe grisea. Mol Gen Genet 251:665–74.

25. Dobinson KF, Harris RE, Hamer JE. 1993. Grasshopper, a long terminal repeat (LTR) retroelement in the phytopathogenic fungus Magnaporthe grisea. Mol Plant Microbe Interact 6:114–26.

26. Kimura M. 1980. A simple method for estimating evolutionary rates of base substitutions through comparative studies of nucleotide sequences. J Mol Evol 16:111–20.

27. Chalopin D, Naville M, Plard F, Galiana D, Volff JN. 2015. Comparative analysis of transposable elements highlights mobilome diversity and evolution in vertebrates. Genome Biol Evol 7:567–80.

28. Peng Z, Oliveira-Garcia E, Lin G, Hu Y, Dalby M, Migeon P, Tang H, Farman M, Cook D, White FF, Valent B, Liu S. 2019. Effector gene reshuffling involves dispensable mini-chromosomes in the wheat blast fungus. PLoS Genet 15:e1008272.

29. Gómez Luciano LB, Tsai IJ, Chuma I, Tosa Y, Chen Y-H, Li J-Y, Li M-Y, Lu M-YJ, Nakayashiki H, Li W-H. 2019. Blast fungal genomes show frequent chromosomal changes, gene gains and losses, and effector gene turnover. Molecular biology and evolution 36:1148–1161.

30. Zhong Z, Chen M, Lin L, Han Y, Bao J, Tang W, Lin L, Lin Y, Somai R, Lu L, Zhang W, Chen J, Hong Y, Chen X, Wang B, Shen W-C, Lu G, Norvienyeku J, Ebbole DJ, Wang Z. 2018. Population genomic analysis of the rice blast fungus reveals specific events associated with expansion of three main clades. The ISME Journal doi:10.1038/s41396-018-0100-6.

31. Zhong Z, Norvienyeku J, Chen M, Bao J, Lin L, Chen L, Lin Y, Wu X, Cai Z, Zhang Q, Lin X, Hong Y, Huang J, Xu L, Zhang H, Chen L, Tang W, Zheng H, Chen X, Wang Y, Lian B, Zhang L, Tang H, Lu G, Ebbole DJ, Wang B, Wang Z. 2016. Directional Selection from Host Plants Is a Major Force Driving Host Specificity in Magnaporthe Species. Sci Rep 6:25591.

32. Lorrain C, Feurtey A, Möller M, Haueisen J, Stukenbrock E. 2021. Dynamics of transposable elements in recently diverged fungal pathogens: lineage-specific transposable element content and efficiency of genome defenses. G3 11:jkab068.

33. Plissonneau C, Hartmann FE, Croll D. 2018. Pangenome analyses of the wheat pathogen Zymoseptoria tritici reveal the structural basis of a highly plastic eukaryotic genome. BMC Biol 16:5.

34. Badet T, Oggenfuss U, Abraham L, McDonald BA, Croll D. 2020. A 19-isolate reference-quality global pangenome for the fungal wheat pathogen Zymoseptoria tritici. BMC Biol 18:12.

35. Torres DE, Oggenfuss U, Croll D, Seidl MF. 2020. Genome evolution in fungal plant pathogens: looking beyond the two-speed genome model. Fungal Biology Reviews.

36. Schuler MA. 2015. P450s in plants, insects, and their fungal pathogens, p 409–449, Cytochrome P450. Springer.

37. Ide M, Ichinose H, Wariishi H. 2012. Molecular identification and functional characterization of cytochrome P450 monooxygenases from the brown-rot basidiomycete Postia placenta. Arch Microbiol 194:243–53.

38. Nazir KH, Ichinose H, Wariishi H. 2011. Construction and application of a functional library of cytochrome P450 monooxygenases from the filamentous fungus Aspergillus oryzae. Appl Environ Microbiol 77:3147–50.

39. Doddapaneni H, Chakraborty R, Yadav JS. 2005. Genome-wide structural and evolutionary analysis of the P450 monooxygenase genes (P450ome) in the white rot fungus Phanerochaete chrysosporium: evidence for gene duplications and extensive gene clustering. BMC Genomics 6:92.

40. Zhang S, Widemann E, Bernard G, Lesot A, Pinot F, Pedrini N, Keyhani NO. 2012. CYP52X1, representing new cytochrome P450 subfamily, displays fatty acid hydroxylase activity and contributes to virulence and growth on insect cuticular substrates in entomopathogenic fungus Beauveria bassiana. J Biol Chem 287:13477–86.

41. Santana M, Queiroz M. 2015. Transposable elements in fungi: a genomic approach. Scientific J Genetics Gen Ther 1:012.

42. Krishnan P, Meile L, Plissonneau C, Ma X, Hartmann FE, Croll D, McDonald BA, Sanchez-Vallet A. 2018. Transposable element insertions shape gene regulation and melanin production in a fungal pathogen of wheat. BMC Biol 16:78.

43. Lewis ZA, Honda S, Khlafallah TK, Jeffress JK, Freitag M, Mohn F, Schubeler D, Selker EU. 2009. Relics of repeat-induced point mutation direct heterochromatin formation in Neurospora crassa. Genome Res 19:427–37.

44. Gladieux P, Condon B, Ravel S, Soanes D, Maciel JLN, Nhani A, Jr., Chen L, Terauchi R, Lebrun MH, Tharreau D, Mitchell T, Pedley KF, Valent B, Talbot NJ, Farman M, Fournier E. 2018. Gene flow between divergent cereal- and grass-specific lineages of the rice blast fungus Magnaporthe oryzae. mBio 9.

45. Gladieux P, Ravel S, Rieux A, Cros-Arteil S, Adreit H, Milazzo J, Thierry M, Fournier E, Terauchi R, Tharreau D. 2018. Coexistence of multiple endemic and pandemic lineages of the rice blast pathogen. mBio 9.

46. Chadha S, Sharma M. 2014. Transposable elements as stress adaptive capacitors induce genomic instability in fungal pathogen Magnaporthe oryzae. PLoS One 9:e94415.

47. Kofler R, Gomez-Sanchez D, Schlotterer C. 2016. PoPoolationTE2: Comparative Population Genomics of Transposable Elements Using Pool-Seq. Mol Biol Evol 33:2759–64.

48. Oggenfuss U, Croll D. 2023. Recent transposable element bursts are associated with the proximity to genes in a fungal plant pathogen. PLoS Pathog 19:e1011130.

49. Wang C, Milgate AW, Solomon PS, McDonald MC. 2021. The identification of a transposon affecting the asexual reproduction of the wheat pathogen Zymoseptoria tritici. Mol Plant Pathol 22:800–816.

50. Fouché S, Badet T, Oggenfuss U, Plissonneau C, Francisco CS, Croll D. 2020. Stress-driven transposable element de-repression dynamics and virulence evolution in a fungal pathogen. Mol Biol Evol 37:221–239.

51. Francis A, Ghosh S, Tyagi K, Prakasam V, Rani M, Singh NP, Pradhan A, Sundaram RM, Priyanka C, Laha GS, Kannan C, Prasad MS, Chattopadhyay D, Jha G. 2023. Evolution of pathogenicity-associated genes in Rhizoctonia solani AG1-IA by genome duplication and transposon-mediated gene function alterations. BMC Biology 21:15.

52. Li W, Wang B, Wu J, Lu G, Hu Y, Zhang X, Zhang Z, Zhao Q, Feng Q, Zhang H, Wang Z, Wang G, Han B, Wang Z, Zhou B. 2009. The Magnaporthe oryzae avirulence gene AvrPiz-t encodes a predicted secreted protein that triggers the immunity in rice mediated by the blast resistance gene Piz-t. Mol Plant Microbe Interact 22:411–20.

53. Zhang S, Wang L, Wu W, He L, Yang X, Pan Q. 2015. Function and evolution of Magnaporthe oryzae avirulence gene AvrPib responding to the rice blast resistance gene Pib. Sci Rep 5:11642.

54. Chen C, Chen M, Hu J, Zhang W, Zhong Z, Jia Y, Allaux L, Fournier E, Tharreau D, Wang G-L. 2014. Sequence variation and recognition specificity of the avirulence gene AvrPiz-t in Magnaporthe oryzae field populations. Fungal Genomics & Biology 4.

55. Miki S, Matsui K, Kito H, Otsuka K, Ashizawa T, Yasuda N, Fukiya S, Sato J, Hirayae K, Fujita Y, Nakajima T, Tomita F, Sone T. 2009. Molecular cloning and characterization of the AVR-Pia locus from a Japanese field isolate of Magnaporthe oryzae. Mol Plant Pathol 10:361–74.

56. Hu ZJ, Huang YY, Lin XY, Feng H, Zhou SX, Xie Y, Liu XX, Liu C, Zhao RM, Zhao WS, Feng CH, Pu M, Ji YP, Hu XH, Li GB, Zhao JH, Zhao ZX, Wang H, Zhang JW, Fan J, Li Y, Peng YL, He M, Li DQ, Huang F, Peng YL, Wang WM. 2022. Loss and Natural Variations of Blast Fungal Avirulence Genes Breakdown Rice Resistance Genes in the Sichuan Basin of China. Front Plant Sci 13:788876.

57. Wu J, Kou Y, Bao J, Li Y, Tang M, Zhu X, Ponaya A, Xiao G, Li J, Li C, Song MY, Cumagun CJ, Deng Q, Lu G, Jeon JS, Naqvi NI, Zhou B. 2015. Comparative genomics identifies the Magnaporthe oryzae avirulence effector AvrPi9 that triggers Pi9-mediated blast resistance in rice. New Phytol 206:1463–75.

58. Xu Z, Wang H. 2007. LTR_FINDER: an efficient tool for the prediction of full-length LTR retrotransposons. Nucleic Acids Res 35:W265–8.

59. Ellinghaus D, Kurtz S, Willhoeft U. 2008. LTRharvest, an efficient and flexible software for de novo detection of LTR retrotransposons. BMC Bioinformatics 9:18.

60. Ou S, Jiang N. 2018. LTR_retriever: A Highly Accurate and Sensitive Program for Identification of Long Terminal Repeat Retrotransposons. Plant Physiol 176:1410–1422.

61. Lu S, Wang J, Chitsaz F, Derbyshire MK, Geer RC, Gonzales NR, Gwadz M, Hurwitz DI, Marchler GH, Song JS, Thanki N, Yamashita RA, Yang M, Zhang D, Zheng C, Lanczycki CJ, Marchler-Bauer A. 2020. CDD/SPARCLE: the conserved domain database in 2020. Nucleic Acids Res 48:D265–D268.

62. Tempel S. 2012. Using and understanding RepeatMasker. Methods Mol Biol 859:29–51.

63. Petersen TN, Brunak S, von Heijne G, Nielsen H. 2011. SignalP 4.0: discriminating signal peptides from transmembrane regions. Nat Methods 8:785–6.

64. Krogh A, Larsson B, von Heijne G, Sonnhammer EL. 2001. Predicting transmembrane protein topology with a hidden Markov model: application to complete genomes. J Mol Biol 305:567–80.

65. Li H, Durbin R. 2009. Fast and accurate short read alignment with Burrows-Wheeler transform. Bioinformatics 25:1754–60.

66. Kofler R, Pandey RV, Schlotterer C. 2011. PoPoolation2: identifying differentiation between populations using sequencing of pooled DNA samples (Pool-Seq). Bioinformatics 27:3435–6.

67. Galili T. 2015. dendextend: an R package for visualizing, adjusting and comparing trees of hierarchical clustering. Bioinformatics 31:3718–20.

68. Danecek P, Auton A, Abecasis G, Albers CA, Banks E, DePristo MA, Handsaker RE, Lunter G, Marth GT, Sherry ST, McVean G, Durbin R, Genomes Project Analysis G. 2011. The variant call format and VCFtools. Bioinformatics 27:2156–8.

69. Purcell S, Neale B, Todd-Brown K, Thomas L, Ferreira MA, Bender D, Maller J, Sklar P, de Bakker PI, Daly MJ, Sham PC. 2007. PLINK: a tool set for whole-genome association and population-based linkage analyses. Am J Hum Genet 81:559–75.

70. Nordberg H, Cantor M, Dusheyko S, Hua S, Poliakov A, Shabalov I, Smirnova T, Grigoriev IV, Dubchak I. 2014. The genome portal of the Department of Energy Joint Genome Institute: 2014 updates. Nucleic Acids Res 42:D26–31.

71. Mistry J, Chuguransky S, Williams L, Qureshi M, Salazar GA, Sonnhammer ELL, Tosatto SCE, Paladin L, Raj S, Richardson LJ, Finn RD, Bateman A. 2021. Pfam: The protein families database in 2021. Nucleic Acids Res 49:D412–D419.

72. Kim D, Paggi JM, Park C, Bennett C, Salzberg SL. 2019. Graph-based genome alignment and genotyping with HISAT2 and HISAT-genotype. Nat Biotechnol 37:907–915.

73. Pertea M, Pertea GM, Antonescu CM, Chang TC, Mendell JT, Salzberg SL. 2015. StringTie enables improved reconstruction of a transcriptome from RNA-seq reads. Nat Biotechnol 33:290–5.

74. Zhang L-M, Chen S-T, Qi M, Cao X-Q, Liang N, Li Q, Tang W, Lu G-D, Zhou J, Yu W-Y, Wang Z-H, Zheng H-K. 2021. The putative elongator complex protein Elp3 is involved in asexual development and pathogenicity by regulating autophagy in the rice blast fungus. Journal of Integrative Agriculture 20:2944–2956.

75. Han Y, Song L, Peng C, Liu X, Liu L, Zhang Y, Wang W, Zhou J, Wang S, Ebbole D, Wang Z, Lu GD. 2019. A Magnaporthe chitinase interacts with a rice jacalin-related lectin to promote host colonization. Plant Physiol 179:1416–1430.

