## Supplementary Tables 1-11 and Supplementary Figures 1-3 for "Transposable elements impact the population divergence of rice blast fungus *Magnaporthe oryzae*": Mo.PopulationTE.manuscript_Apr_2023.supplementary figure_v3.docx

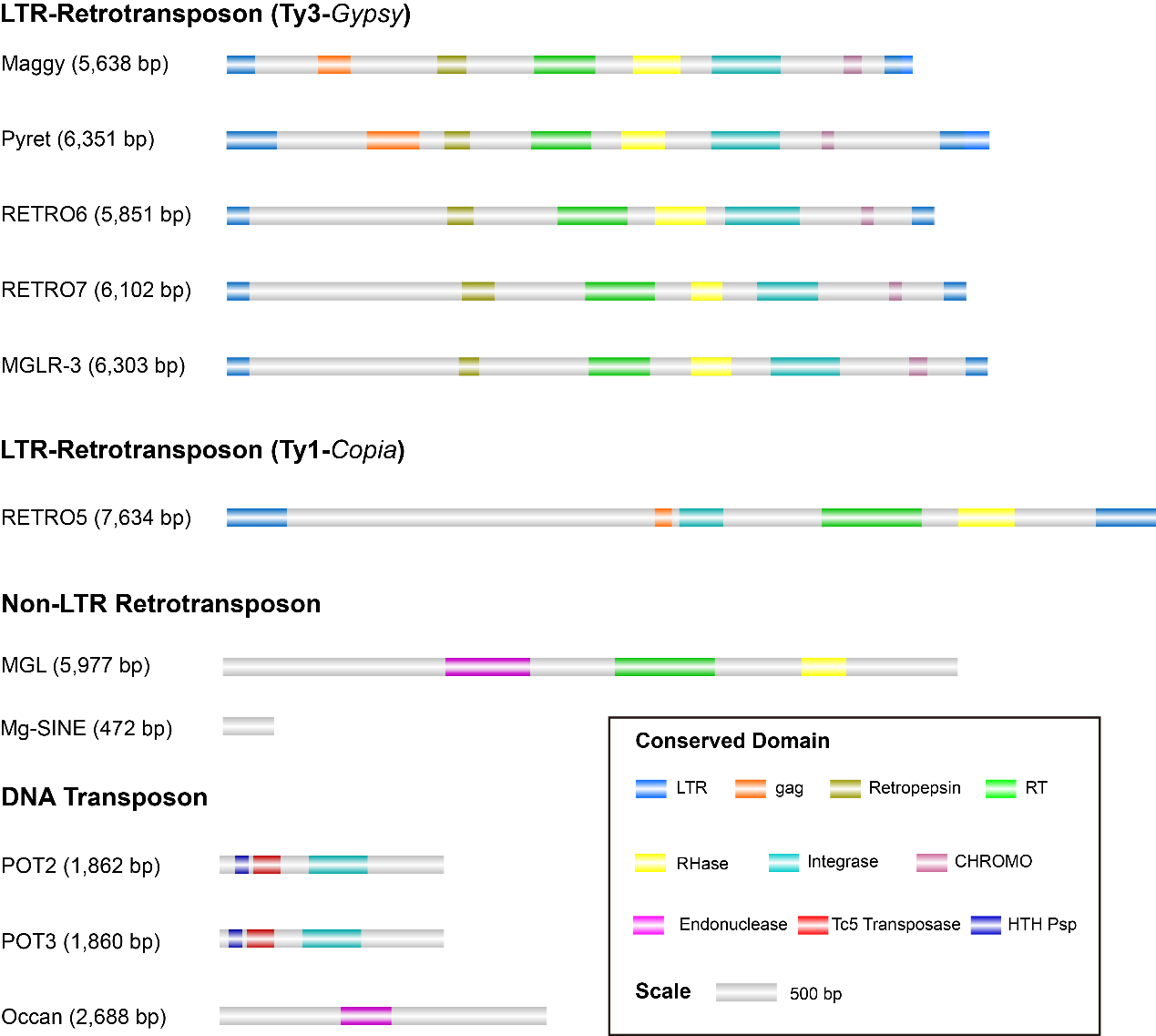

**FIG S1** Schematic of the 11 most abundant TE families on the genome of the *M. oryzae* rice isolate. The conserved domains in the relative location of the TE consensus were highlighted using different colors.

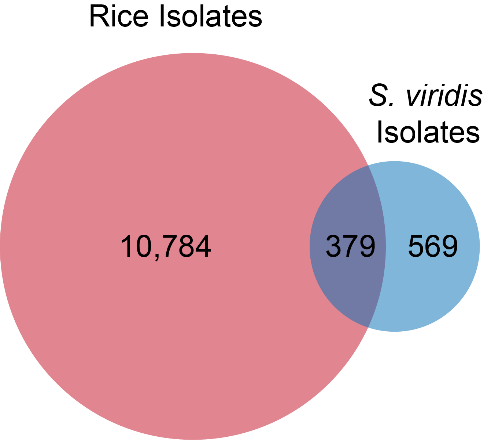

**FIG S2** The venn chart compares the TE insertion loci of 90 *M. oryzae* rice isolates and two *S. viridis* isolates.

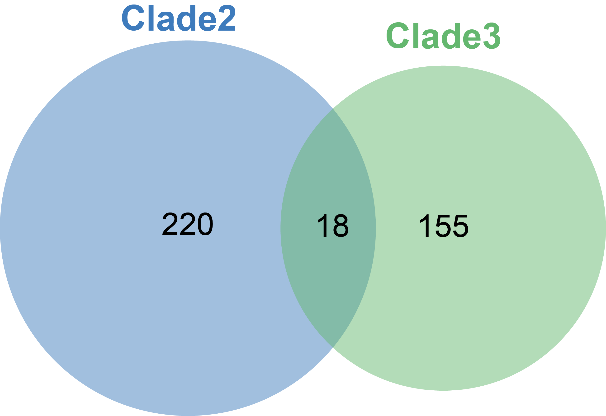

**FIG S3** The venn chart displays the intersection of clade2-specific and clade3-specific TE-associated genes.
