## Supplementary Tables 1-11 and Supplementary Figures 1-3 for "Transposable elements impact the population divergence of rice blast fungus *Magnaporthe oryzae*": Mo.PopulationTE.manuscript_Apr_2023.supplementary table R2.docx

**TABLE S2** The proportions of 11 TE families on the genomes of a *M. oryzae* wheat isolate and three grass isolates.

| **TE Type** | **Wheat Isolate** |  | **Grass Isolates** | | |
| --- | --- | --- | --- | --- | --- |
|  | **B71** |  | **MZ5-1-6** | **NI907** | ***NI919*** |
| **MGLR-3** | 0.30% |  | 0.10% | 0.12% | *0.64%* |
| **Maggy** | 0.00% |  | 0.00% | 0.00% | *0.00%* |
| **Pyret** | 1.77% |  | 1.89% | 1.67% | *2.60%* |
| **RETRO6** | 0.42% |  | 0.29% | 0.07% | *0.95%* |
| **RETRO7** | 0.64% |  | 0.24% | 0.68% | *0.27%* |
| **RETRO5** | 0.46% |  | 0.43% | 0.22% | 0.03% |
| **MGL** | 0.69% |  | 0.23% | 0.62% | 0.12% |
| **Mg-SINE** | 0.02% |  | 0.01% | 0.01% | 0.00% |
| **POT2** | 0.29% |  | 0.64% | 0.51% | 0.05% |
| **POT3** | 0.25% |  | 0.11% | 0.14% | 0.53% |
| **Occan** | 0.16% |  | 0.05% | 0.11% | 0.02% |
| **Total** | 5.00% |  | 3.99% | 4.15% | 5.21% |

| **TABLE S3** Validation of 17 TE insertion sites by PacBio. | | | | | | | | |
| --- | --- | --- | --- | --- | --- | --- | --- | --- |
| **70-15** | | | | | **81278** | | **Guy11** | |
| **Chromosome** | **TE insertion site** | **Isolates containing TE insertion** | **TE type** | **Associated Gene** | **Exist in 81278** | **Genomic location** | **Exist in Guy11** | **Genomic location** |
| Supercontig 8.1 | 1,045,432 | FJ81278 | Occan | MGG_16070 | **Y** | MQOO01000006.1:76294-78772 | N | - |
| Supercontig 8.2 | 337,923 | Guy11 | POT2 | MGG_10466 | N | - | **Y** | MQOP01000015.1:712,892-714,750 |
| Supercontig 8.2 | 5,189,603 | Guy11 | POT2 | MGG_01879 | N | - | **Y** | MQOP01000031.1:17,249-19,110 |
| Supercontig 8.2 | 7,361,446 | Guy11 | POT2 | MGG_15908 | N | - | **Y** | MQOP01000001.1:474,327-476,187 |
| Supercontig 8.3 | 124,514 | Guy11 | MGL | MGG_15185 | N | - | **Y** | MQOP01000021.1:88,201-88,774 |
| Supercontig 8.3 | 219,010 | Guy11 | POT3 | MGG_16593 | N | - | **Y** | MQOP01000021.1:182,069-183,929 |
| Supercontig 8.4 | 4,455,773 | FJ81278 | POT2 | MGG_06234 | **Y** | MQOO01000003.1:3,680,026-3,681,886 | N | - |
| Supercontig 8.4 | 4,893,019 | Guy11 | POT3 | MGG_17244 | N | - | **Y** | MQOP01000019.1:193,259-195,119 |
| Supercontig 8.5 | 243,556 | Guy11 | POT2 | MGG_10796 | N | - | **Y** | MQOP01000014.1:57,995-59,856 |
| Supercontig 8.5 | 2,041,379 | Guy11/FJ81278 | POT2 | MGG_00677 | **Y** | MQOO01000007.1:280,227-282,087 | **Y** | MQOP01000015.1:712,892-714,750 |
| Supercontig 8.6 | 86,055 | FJ81278 | POT2 | MGG_08796 | **Y** | MQOO01000002.1:2,367,307-2,369,168 | N | - |
| Supercontig 8.6 | 562,159 | Guy11 | POT3 | MGG_17590 | N | - | **Y** | MQOP01000011.1:615,793-617,653 |
| Supercontig 8.6 | 588,503 | Guy11 | POT2 | MGG_01943 | N | - | **Y** | MQOP01000014.1:57,995-59,856 |
| Supercontig 8.6 | 3,326,207 | FJ81278 | POT2 | MGG_09425 | **Y** | MQOO01000012.1:251,995-253,855 | N | - |
| Supercontig 8.7 | 2,411,904 | Guy11 | POT2 | MGG_02546 | N | - | **Y** | MQOP01000022.1:52,970-54,830 |
| Supercontig 8.7 | 2,907,029 | Guy11/FJ81278 | POT2 | MGG_10559 | **Y** | MQOO01000034.1:16000-17864 | **Y** | MQOP01000012.1:421,276-423,138 |
| Supercontig 8.8 | 83,870 | FJ81278 | POT2 | MGG_15046 | **Y** | MQOO01000008.1:1,939,603-1,941,464 | N | - |

**TABLE S4** Validation of 17 TE insertion sites by PCR amplification.

| **Gene** | **Chr** | **Position** | **TE** | **Isolates** | **Predicted by Reads** | **Validated by PCR** |
| --- | --- | --- | --- | --- | --- | --- |
| MGG_16070 | Supercontig_8.1 | 1,045,432 | Occan | FJ81278 | Y | Y |
| MGG_10466 | Supercontig_8.2 | 337,923 | POT2 | Guy11 | Y | Y |
| MGG_01879 | Supercontig_8.2 | 5,189,603 | POT2 | Guy11 | Y | Y |
| MGG_15908 | Supercontig_8.2 | 7,361,446 | POT2 | Guy11 | Y | Y |
| MGG_15185 | Supercontig_8.3 | 124,514 | MGL | Guy11 | Y | Y |
| MGG_16593 | Supercontig_8.3 | 219,010 | POT3 | Guy11 | Y | Y |
| MGG_06234 | Supercontig_8.4 | 4,455,773 | POT2 | FJ81278 | Y | Y |
| MGG_17244 | Supercontig_8.4 | 4,893,019 | POT3 | Guy11 | Y | Y |
| MGG_10796 | Supercontig_8.5 | 243,556 | POT2 | Guy11 | Y | Y |
| MGG_00677 | Supercontig_8.5 | 2,041,379 | POT2 | Guy11/FJ81278 | Y | Y |
| MGG_08796 | Supercontig_8.6 | 86,055 | POT2 | FJ81278 | Y | Y |
| MGG_17590 | Supercontig_8.6 | 562,159 | POT3 | Guy11 | Y | Y |
| MGG_01943 | Supercontig_8.6 | 588,503 | POT2 | Guy11 | Y | Y |
| MGG_09425 | Supercontig_8.6 | 3,326,207 | POT2 | FJ81278 | Y | Y |
| MGG_02546 | Supercontig_8.7 | 2,411,904 | POT2 | Guy11 | Y | Y |
| MGG_10559 | Supercontig_8.7 | 2,907,029 | POT2 | Guy11/FJ81278 | Y | Y |
| MGG_15046 | Supercontig_8.8 | 83,870 | POT2 | FJ81278 | Y | Y |

**TABLE S5** Insertion preference of various TEs in different genomic regions.

| **TE Family** | **Genomic Region** | **Number in Genomic Region** | **Total Number** | **FC** | **P-value (adjusted)** |
| --- | --- | --- | --- | --- | --- |
| POT2 | Flanking | 2,462 | 4,027 | 1.10 | 2.53E-18 |
| SINE | Intragenic | 458 | 1,009 | 1.39 | 1.90E-17 |
| MGL | Intragenic | 620 | 1,487 | 1.27 | 5.21E-14 |
| Maggy | Intergenic | 242 | 692 | 1.51 | 1.59E-12 |
| MGLR-3 | Intergenic | 83 | 210 | 1.71 | 4.02E-07 |
| RETRO6 | Intragenic | 201 | 492 | 1.25 | 3.80E-04 |
| POT3 | Intergenic | 266 | 945 | 1.22 | 4.71E-04 |
| RETRO7 | Intergenic | 160 | 539 | 1.28 | 7.43E-04 |
| RETRO5 | Intergenic | 97 | 306 | 1.37 | 1.10E-03 |
| Occan | Intergenic | 86 | 282 | 1.32 | 7.20E-03 |
| Pyret | Intergenic | 311 | 1,174 | 1.15 | 7.20E-03 |

#FC: fold change, counted based on the formula: (number of specific TEs in region / number of all TEs in the region) / (number of specific TEs / number of all TEs).

**TABLE S6** Enrichment analysis of TEs in the promoter of secreted proteins.

| **Region** | **TE Type** | **Associated with Secreted Proteins** | **Associated with All Genes** | **FC** | **P-value**  **(adjusted)** |
| --- | --- | --- | --- | --- | --- |
| **Upstream** | POT2 | 257 | 1,138 | 1.15 | 5.28E-03 |
|  | POT3 | 88 | 305 | 1.47 | 6.50E-04 |

#FC: fold change, which is defined as the proportion of specific TEs correlated to secreted proteins accounting for all TEs correlated to secreted proteins divided by the proportion of specific TEs correlated to all genes accounting for all TEs correlated to all genes.

**TABLE S7** Identity of TE insertion sites between intra- or inter-clade isolates.

|  | **Clade1** | **Clade2** | **Clade3** |
| --- | --- | --- | --- |
| **Clade1** | 34.16% | 29.22% | 29.09% |
| **Clade2** | 29.22% | 59.38% | 27.71% |
| **Clade3** | 29.09% | 27.71% | 55.34% |

**TABLE S8** Enrichment analysis of the clade2- and clade3-specific TE-associated genes using conserved domains.

| **Clade** | **Pfam**  **Annotation** | **Clade-specific genes** | **All genes** | **Fold Change** | **P-value**  **(adjusted)** |
| --- | --- | --- | --- | --- | --- |
| **Clade2** | PF00067:  Cytochrome P450 | 8 | 79 | 6.47 | 2.12E-03 |
| **Clade3** | PF00067:  Cytochrome P450 | 9 | 67 | 8.59 | 1.21E-04 |

**TABLE S9** GEO information of 16 RNA seq samples.

| **Sample Name** | **Reads Name** | **GEO ID** |
| --- | --- | --- |
| 13FM-24-1_rep1 | 13FM-24-1_FRAS210075990-1 | GSM6210796 |
| 13FM-24-1_rep2 | 13FM-24-1_FRAS210075990-2 | GSM6210797 |
| AV1-1-1_rep1 | AV1-1-1_FRAS210075995-1 | GSM6210798 |
| AV1-1-1_rep2 | AV1-1-1_FRAS210075995-2 | GSM6210799 |
| BJ08-8_rep1 | BJ08-8_FRAS210076005-1 | GSM6210800 |
| BJ08-8_rep2 | BJ08-8_FRAS210076005-2 | GSM6210801 |
| DB11-621_rep1 | DB11-621_FRAS210076004-1 | GSM6210802 |
| DB11-621_rep2 | DB11-621_FRAS210076004-2 | GSM6210803 |
| FJ72AC7-77_rep1 | FJ72AC7-77_FRAS210075997-1 | GSM6210804 |
| FJ72AC7-77_rep2 | FJ72AC7-77_FRAS210075997-2 | GSM6210805 |
| FJ78-JJ_rep1 | FJ78-JJ_FRAS210076001-1 | GSM6210806 |
| FJ78-JJ_rep2 | FJ78-JJ_FRAS210076001-2 | GSM6210807 |
| FJ81278_rep1 | FJ81278_FRAS210076000-2 | GSM6210808 |
| FJ81278_rep2 | FJ81278_FRAS210076000-3 | GSM6210809 |
| FJ8773-19_rep1 | FJ8773-19_FRAS210075992-1 | GSM6210810 |
| FJ8773-19_rep2 | FJ8773-19_FRAS210075992-2 | GSM6210811 |
| FJ98099_rep1 | FJ98099_FRAS210075996-1 | GSM6210812 |
| FJ98099_rep2 | FJ98099_FRAS210075996-2 | GSM6210813 |
| FJSH0703_rep1 | FJSH0703_FRAS210075998-1 | GSM6210814 |
| FJSH0703_rep2 | FJSH0703_FRAS210075998-2 | GSM6210815 |
| GD06-53_rep1 | GD06-53_FRAS210075999-1 | GSM6210816 |
| GD06-53_rep2 | GD06-53_FRAS210075999-2 | GSM6210817 |
| Nich-2-3-2_rep1 | Nich-2-3-2_FRAS210075991-1 | GSM6210818 |
| Nich-2-3-2_rep2 | Nich-2-3-2_FRAS210075991-2 | GSM6210819 |
| Sar-2-20-1_rep1 | Sar-2-20-1_FRAS210075994-1 | GSM6210820 |
| Sar-2-20-1_rep2 | Sar-2-20-1_FRAS210075994-3 | GSM6210821 |
| TW-PT1-1_rep1 | TW-PT1-1_FRAS210076003-1 | GSM6210822 |
| TW-PT1-1_rep2 | TW-PT1-1_FRAS210076003-2 | GSM6210823 |
| YN072313_rep1 | YN072313_FRAS210075993-1 | GSM6210824 |
| YN072313_rep2 | YN072313_FRAS210075993-2 | GSM6210825 |
| ZJ00-10_rep1 | ZJ00-10_FRAS210076002-1 | GSM6210826 |
| ZJ00-10_rep2 | ZJ00-10_FRAS210076002-2 | GSM6210827 |
